# A sensitivity-consistency trade-off in memory formation regulated by dentate gyrus inhibition

**DOI:** 10.1101/2025.07.28.667209

**Authors:** Elena Pérez-Montoyo, José María Caramés, Raquel Garcia-Hernandez, Santiago Canals, Encarni Marcos

## Abstract

A core computational challenge for memory systems is the need to balance two opposing demands: (1) sensitivity to subtle input differences for flexible encoding, and (2) consistency to preserve stable, interference-resistant representations. How the brain dynamically regulates this trade-off remains unclear. Here, we show that modulating parvalbumin-expressing (PV^+^) interneurons in the dentate gyrus regulates this balance by shifting hippocampal computation between sensitivity and consistency regimes. Combining cell-type–specific pharmacogenetics, behavioral assays, and computational modeling, we found that reducing PV^+^-mediated inhibition during encoding enhanced the subsequent discrimination of similar inputs but it increased vulnerability to interference. In contrast, increased inhibition stabilized memory representations at the cost of discriminability. Together, the results establish PV^+^ interneurons as key modulators of hippocampal memory processing, enabling the system to prioritize either flexible updating or robust retention based on inhibitory tone at the time of encoding. This sensitivity–consistency continuum may reflect a fundamental organizing principle of memory computation.

## Introduction

How the brain flexibly encodes new experiences while preserving stable memories of the past remains a central challenge in system neuroscience ^1^. Episodic memory formation demands a balance between sensitivity - the ability to detect and encode subtle but behaviorally relevant differences across similar experiences - and consistency - the capacity to preserve stable, interference-resistant memory traces over time. Yet, the mechanisms that support this dynamic trade-off remain poorly understood. Here, we address this problem by manipulating a key inhibitory component of the hippocampal circuit and examining how it influences memory performance under different computational demands.

The hippocampus plays a central role in encoding episodic experiences. Within it, the dentate gyrus (DG) has been implicated in disambiguating similar events through pattern separation - the transformation of similar inputs into distinct neural representations ^2, 3^. A key feature supporting pattern separation in the DG is the sparse activation of granule cells (GCs), which is a feature thought to enhance the distinctiveness of memory representations ^4, 5^. This sparsity is enhanced, in part, by fast-spiking, parvalbumin-expressing (PV^+^) interneurons. PV^+^ cells receive excitatory input from both the entorhinal cortex and the ipsi- and contralateral hippocampus ^6^, and implement competitive dynamics that restrict GC activity through a winner-take-all mechanism ^7-10^. Through this circuit, PV^+^ interneurons are thought to promote sparse, decorrelated activity in the DG that reduces overlap between similar memories.

Beyond enforcing sparsity, PV^+^ interneurons modulate inhibition in a dynamic, experience-dependent manner. This form of regulation is increasingly recognized as essential for learning and memory ^11-15^. Periods of transient disinhibition, often achieved through reduced PV^+^ activity, have been causally linked to the encoding and expression of memory ^11, 12^. Moreover, memory formation has been associated with plastic changes within the PV⁺ interneuron network, suggesting that flexible, experience-dependent modulation of the inhibitory tone may be a fundamental mechanism for memory formation ^16^.

However, the above findings raise a fundamental computational conundrum: the same disinhibition that enhances the encoding of new experiences would also undermine the distinctiveness of stored representations. In other words, how does the hippocampal network resolve the competing demands of encoding flexibility and memory stability? To address this question, we combined cell-type-specific pharmacogenetics, behavioral assays and computational modeling. We systematically manipulated PV^+^ interneuron activity in the DG during the encoding phase of spatial memory tasks, and assessed its impact on later spatial discrimination and memory stability across varying interference demands. We complemented this with a biologically inspired computational model of cortico-hippocampal circuits that simulates novelty detection via novelty-driven decision making. Our findings reveal that PV^+^ interneurons may serve as a dynamic control mechanism that shifts hippocampal computation between sensitivity- and consistency-dominant regimes. This trade-off, governed by local inhibitory dynamics, may reflect a core organizing principle of hippocampal memory computation.

## Results

We manipulated the activity of PV^+^ interneuron in the DG pharmacogenetically to examine its impact on memory performance. Prior to behavioral testing, we validated the specificity and efficacy of this manipulation using *in vivo* electrophysiological recordings and *postmortem* immunostaining of PV^+^ cells and c-Fos quantification.

### DG PV^+^ cells regulate granule cells output bidirectionally

We regulated PV-cell activity in the DG with Designer Receptors Exclusively Activated by Designer Drugs (DREADDs) ^17^. These receptors allow gain- or loss-of-function manipulations of PV-cell firing, depending on the subtype of receptor expressed, without imposing an external pattern of activation nor completely blocking spiking. We injected stereotaxically adeno-associated virus 5 (AAV5)-human synapsin1 (hSyn)-DIO-hM4D(Gi)-mCherry into the hilus of the DG of PV-cre transgenic mice (129P2-Pvalbtm1(cre)Arbr/J) to express the inhibitory hM4D(Gi) receptor in dentate PV^+^ interneurons (Fig. 1a; referred from now as PV-Gi animals). Conversely, for the activation of PV-cells we injected the AAV5-hSyn-DIO-hM3D(Gq)-mCherry virus to express the excitatory hM3D(Gq) receptor (Fig. 1a; referred to as PV-Gq animals). The proportion of PV^+^ interneurons infected in the DG was 92.8 ± 3.8 % and 91.2 ± 3.5 % for hM4D(Gi) or hM3D(Gq) viruses, respectively (Fig. S1a-c).

**Figure 1.**
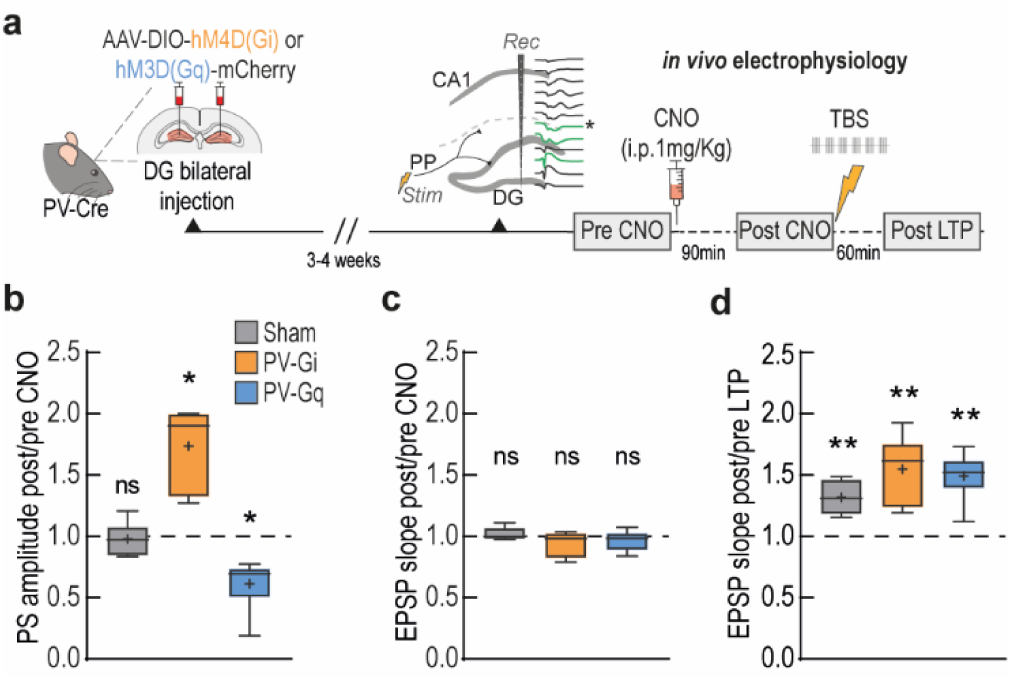
Modulation of PV-cell activity in the DG controls GC output preserving synaptic activity and plasticity. **(a)** Experimental protocol for *in vivo* electrophysiology. PV cells were manipulated using adeno-associated viruses (AAV) carrying designer receptors exclusively activated by designer drugs (DREADDs), allowing inhibition (hM4D(Gi)) or activation (hM3D(Gq)) of these cells. After allowing 3-4 weeks for viral expression, electrophysiological responses in the DG upon perforant path stimulation were recorded before and after CNO injection and LTP induction. **(b-c)** Ratio of the population spike (PS) **(b)** and the excitatory postsynaptic potential (EPSP) **(c)** evoked by perforant path stimulation between pre- and post-CNO periods. **(d)** LTP of the EPSP expressed as the ratio of the EPSP post-CNO administration before and after LTP induction. All statistical values are detailed in Table S1. * = p<0.05, ** = p<0.01.

Electrophysiological recordings were performed *in vivo* before and 90 min after Clozapine-N-Oxide (CNO) administration (i.p. 1 mg/Kg) and again 60 min following the induction of long-term potentiation (LTP). In PV-Gi animals, evoked electrophysiological potentials in the DG following stimulation of the perforant pathway, the main entorhinal cortex input to the hippocampus, showed a significant increase in the amplitude of the population spike (PS) after CNO (i.p. 1 mg/Kg) administration (Fig. 1b), reflecting the facilitated firing of GC. Conversely, decreased PS amplitude in the DG (Fig. 1b) was recorded in PV-Gq animals after CNO administration, indicative of the hindered GC firing. Importantly, the excitatory postsynaptic potentials (EPSPs), reflecting the synaptic responses and dendritic integration in GC, were unchanged in all groups (Fig. 1c). The integrity of the dendritic function was further demonstrated in synaptic plasticity experiments in which the induction of LTP by high-frequency stimulation of the perforant pathway increased the EPSP in all groups (Fig. 1d).

### DG PV^+^ cells modulation alters spatial discrimination

We then investigated whether the modulation of GC’s output, while preserving intact synaptic and dendritic function, had any impact on memory formation. To this end, we used the hippocampal-dependent novel object location task (NOL) ^18, 19^. In this task (Fig. 2a), mice first explore a context (habituation) and are then reintroduced to the same context containing two identical objects during an exploration session (familiarization). After 24 h, they are re-exposed to the same context, but with one object displaced (test). If spatial memory is intact, animals show a preference for the object that occupies a novel position (see Methods and Fig. 2a).

**Figure 2.**
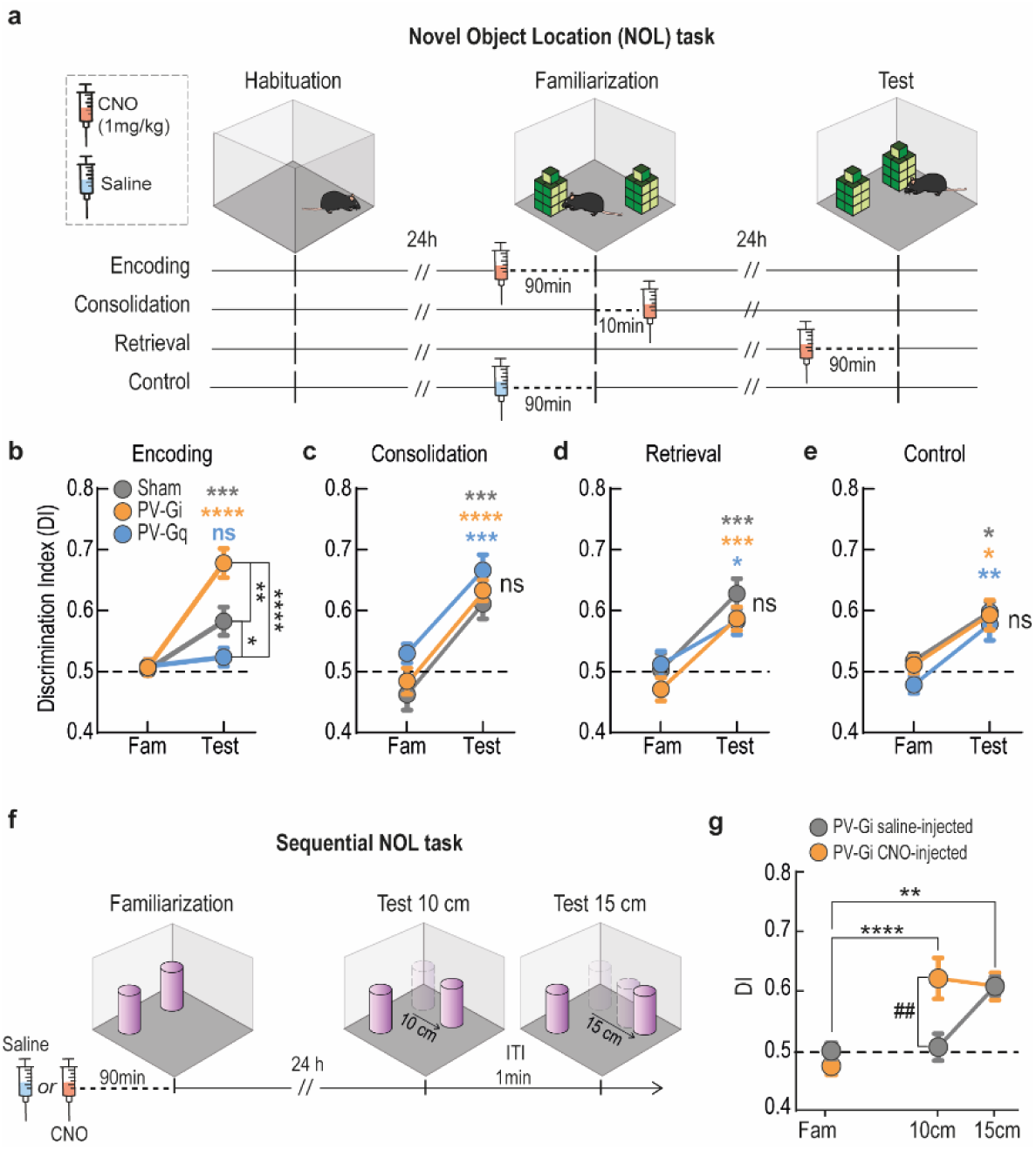
Memory encoding under modulation of PV^+^ cells activity. **(a)** NOL protocols used to modulate PV-cell activity in different task phases. For the control experiment, all groups were saline-injected (no PV-modulation). **(b-e)** Performance in the NOL task with interventions in encoding **(b)**, consolidation **(c)** or retrieval **(d)**, and the saline control **(e)**. Values higher than 0.5 denote preference towards the moved object (mean ± SEM). * within group (Fam. vs. test), # between groups (Sham vs. PV-Gi vs. PV-Gq). (**f)** Sequential NOL task. The object is first displaced 10 cm, and then 5 cm more to the final position at 15 cm from the initial location. CNO (orange) or saline (grey) were injected in the familiarization phase. **(g)** Enhanced novelty detection. Exploration index for each experimental condition (familiarization and 10 and 15 cm object displacements). The same animals were used for both conditions (with saline or CNO administration, respectively). Data represent mean ± SEM. * within group comparison (Fam. vs. test), # CNO vs. saline. All statistical values are detailed in Table S1. * = p<0.05, ** = p<0.01, *** = p<0.001.

To assess the role of DG PV^+^ interneurons during distinct stages of memory processing, we administered CNO (i.p. 1 mg/Kg) either 90 min before encoding, 10 min after encoding (consolidation), or 90 min before retrieval (Fig. 2a). Behavioral performance, measured as a discrimination index (DI; see Methods), was significantly influenced by PV^+^ cell activity modulation during memory encoding (Fig. 2b), but not during consolidation (Fig. 2c) or retrieval (Fig. 2d). Reducing perisomatic inhibition (PV-Gi) during encoding improved memory performance 24 h later (Fig 2b), whereas increasing PV^+^ interneuron activity (PV-Gq) impaired it (Fig. 2b). These memory-related effects were not attributable to differences in locomotor activity or anxiety-like behavior, as assessed by open-field exploration and the elevated plus maze (Fig. S3b-e). As a critical control, the NOL task was repeated in the same group of animals, substituting CNO with its vehicle (saline) during encoding. Under these conditions, all groups showed comparable memory performance (Fig. 2e), confirming that the observed effects were specific to DREADDs activation.

To validate the effectiveness of DREADDs-mediated modulation of PV^+^ interneurons during behavior, we stained c-Fos protein and quantified c-Fos/PV co-labeling in the DG following the familiarization phase of the NOL task (Fig. S2a). This analysis revealed a significant reduction in the number of PV^+^ cells expressing c-Fos in PV-Gi animals and a significant increase in PV-Gq animals following memory encoding (Fig. S2b), consistent with effective pharmacogenetic modulation. Additionally, the overall sensitivity of the task to activate cell assemblies in the DG was confirmed by a higher number of c-Fos^+^ cells after NOL exploration compared to home-cage controls (Fig. S2c-d).

We next asked whether the improved memory performance observed with reduced PV^+^ cell activity came at the expense of spatial discrimination, a function thought to benefit from the sparse activation of GCs ^4, 5^. To address this, we used a sequential NOL task (see Methods), in which animals were exposed to progressively more challenging spatial changes: an initial test with a subtle displacement (10 cm), followed by a second test with a larger displacement (15 cm). PV^+^ cell activity reduction enhanced discrimination sensitivity, particularly under the most challenging condition (Fig. 2f-g). When the displacement was small (10 cm), PV-Gi animals injected with saline failed to detect the change, while the same animals under CNO reliably identified it, indicating increased sensitivity to subtle spatial changes. When the displacement was increased to 15 cm, under both conditions, animals successfully detected the change, suggesting that reduced perisomatic inhibition enhances performance under high-demand conditions.

Together, these results suggest that perisomatic inhibition via PV^+^ interneurons influences the precision of spatial encoding, with reduced inhibition enhancing discrimination. In line with this interpretation, the impaired performance observed in PV-Gq animals likely reflects a reduction in discrimination capacity rather than a general deficit in spatial memory encoding. We next explored this issue. Furthermore, these findings raised an important question about the broader consequences of modulating inhibition: if a lower inhibitory tone improves performance, why doesn’t the system operate in that state by default?

### Computational insights into inhibition-mediated memory dynamics

We developed a cortico-hippocampal neural network designed to emulate an animal’s decision-making process during exploration of a familiar environment, where objects may be encountered in familiar or novel locations. In this model, the level of novelty in a "sensory" input pattern determines whether the model generates a response to explore the novel object location. The model consists of two modules (Fig. 3a), designed to capture key aspects of hippocampal and prefrontal cortex function: the hippocampal-related module implements Hebbian learning and discrimination, while the prefrontal-related module implements decision making.

**Figure 3.**
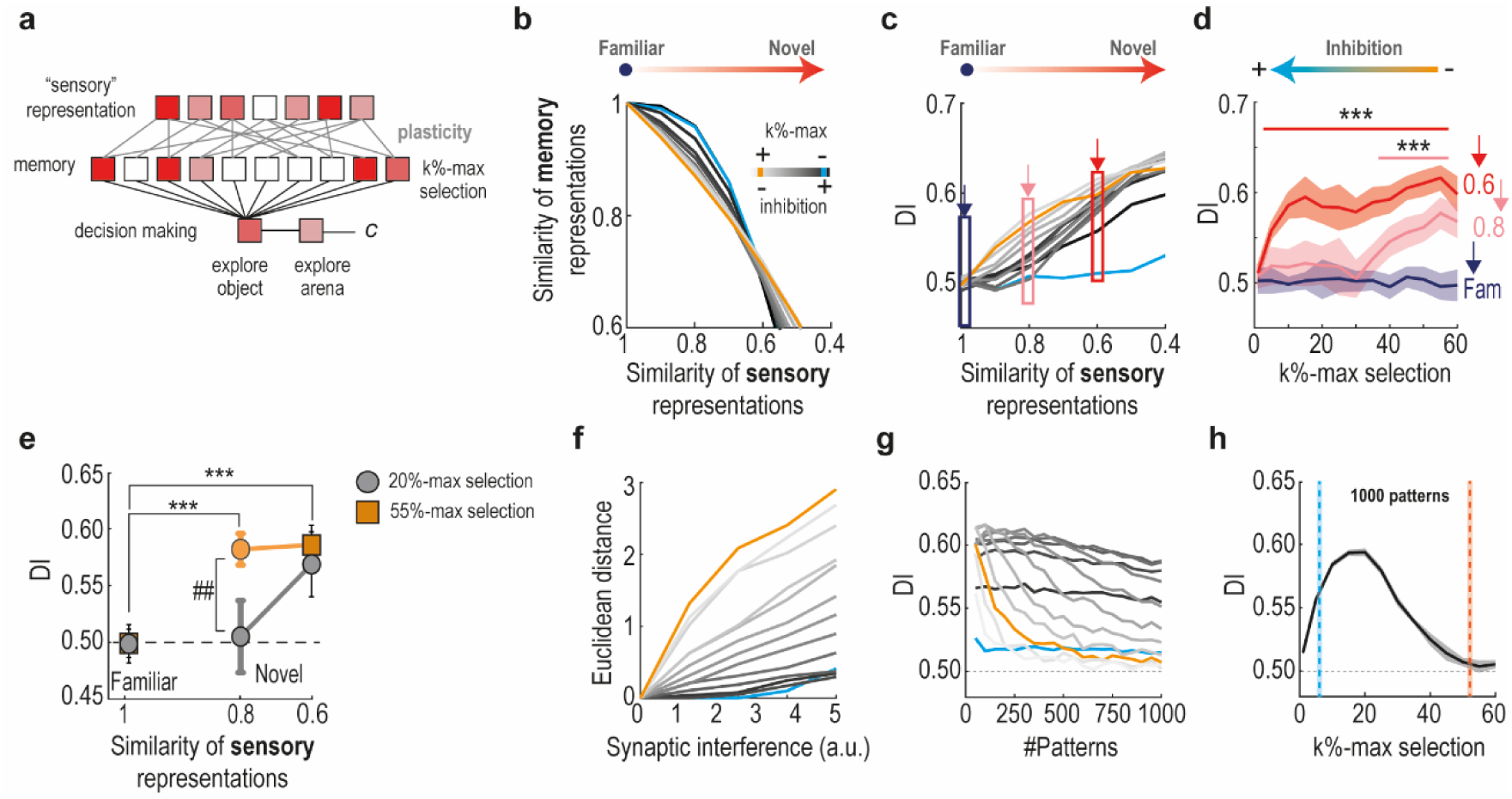
Simulation results show a sensitivity-consistency trade-off in memory formation regulated by the inhibitory tone. **(a)** Scheme of the computational model. It is composed of a “sensory” input layer, a memory layer and decision-making layer, connected with certain probability. Sensory patterns are learned through synaptic plasticity, which modifies the connection strength between the input and memory layers. The memory representation of the input pattern is projected onto the decision-making module, which determines whether to explore the object or the arena. Inhibition levels are modulated using a k%-max selection algorithm, where k indicates the level of inhibition. **(b)** Similarity between memory representations, measured as the correlation between activity patterns evoked by a gradually modified version of a learned sensory input, across different inhibition levels. The blue and orange curves illustrate examples of high and low inhibition levels, respectively. **(c)** Discrimination Index (DI) values for the same modified inputs, showing how pattern separation varies with inhibitory strength. **(d)** DI across different inhibition levels (k%-max selection), corresponding to specific input pattern modifications indicated by the dark blue, dark red and light red arrows in (c) (***=p<0.0001, Two-way ANOVA; mean ± STD). **(e)** DI for specific input similarity values at two levels of inhibition. An input similarity (correlation) of 1 simulates familiar inputs, while values of 0.8 and 0.6 simulate increasing novelty (* within group, # between groups, Two-away ANOVA; mean ± STD). **(f)** Euclidean distance between the memory representation of a learned input pattern and the representation generated by the same pattern after introducing synaptic weights interference. **(g)** DI across varying numbers of learned patterns and different inhibition levels. **(h)** DI at different levels of inhibition after the network learns 1000 sensory input patterns.

To simulate the inhibitory control exerted by PV^+^ interneurons in the DG, we applied a k%-max winner-take-all rule in the hippocampal-related module ^9^ (see Methods). The k% parameter determines the neurons that are allowed to fire: higher k% values correspond to lower inhibition, facilitating neuronal activation, while lower k% values indicate stronger inhibition, limiting neuronal activation. For each inhibition level, the model was trained with a predefined set of “sensory” input patterns, representing activity from a third neuronal layer. It was then tested with gradually modified versions of those patterns to simulate increasing novelty (see Methods). The network’s ability to maintain stable memory representations depended critically on inhibition (Fig. 3b). Under high inhibition (low k%), representations in the hippocampal layer remained stable across a wider range of input pattern changes, suggesting that stability was preserved at the cost of reduced sensitivity. In contrast, under low inhibition (high k%), even small changes in input produced substantial alterations in output representations, indicating increased sensitivity at the cost of stability. This trade-off was reflected in the steeper decline of output similarity (measured as correlation) for similar inputs under low inhibition (Fig. 3b).

To connect these representational changes to behavior, we projected the hippocampal output to the prefrontal-related module, where the model simulated exploration decisions based on the familiarity of incoming sensory patterns. We computed a DI to quantify the model’s tendency to explore objects versus the arena, depending on input novelty (see Methods). As suggested by the hippocampal patterns, the relationship between input similarity and discrimination depended strongly on inhibition level: DI values were higher under low inhibition, especially for subtly modified inputs, reflecting greater sensitivity to change (Fig. 3c). This sensitivity was further quantified by comparing DI across inhibition levels for specific novelty conditions (Fig. 3d), revealing a marked enhancement in novelty detection at low inhibition (high k%) when input similarity was high (0.8). At lower input similarity (0.6), discrimination was less sensitive to disinhibition, and higher inhibition levels still permitted successful discrimination (Fig. 3d). These simulations closely mirrored our experimental findings. When we selected two inhibition levels to reflect control and PV-Gi conditions (intermediate and low inhibition, respectively), the model reproduced the discrimination effect observed in the sequential NOL task (compare Fig. 2g and 3e).

These results suggest that reduced inhibition improves discrimination by amplifying the networks’ responsiveness to subtle input changes. However, this increased sensitivity may come at a cost. In biologically realistic networks subject to noise and ongoing plasticity, excessive sensitivity could destabilize memory representations. To examine this possibility, we introduced synaptic interference into the model by allowing stochastic fluctuations in synaptic weights. This manipulation caused greater disruption under low inhibition, as reflected by an increased Euclidean distance between the memory representations evoked by the same input before and after interference (Fig. 3f). In contrast, high inhibition preserved more stable representations across conditions. These findings highlight the critical role of inhibition in preserving memory stability and raise an important question: does reduced inhibition compromise memory fidelity by increasing susceptibility to synaptic plasticity-induced changes?

To investigate this, we systematically increased the number of learned patterns in the network and calculated the DI for each condition. Under low inhibition, novelty-driven exploration declined sharply with increased memory load, eventually reaching chance levels (DI ≈ 0.5), indicating that familiar and novel inputs could no longer be distinguished (Fig. 3g). In contrast, higher levels of inhibition preserved discriminability across memory loads. However, this came at the cost of sensitivity: very high inhibition levels were also associated with reduced DI, likely because excessive inhibitory tone produced memory representations that were robust to input changes. As a result, the network’s ability to form sufficiently differentiated representations for similar, but unfamiliar, patterns was diminished, leading to reduced exploration of novel stimuli. These effects are further illustrated in Fig. 3h, which shows DI across inhibition levels for a fixed, high memory load. The resulting bell-shaped curve indicates that both low and high inhibition levels impair discrimination in this regime. Together, these results reveal a crucial trade-off: while low inhibition supports finer discrimination, it compromises memory stability by increasing susceptibility to interference. High inhibition, conversely, protects against interference, but limits the network’s sensitivity to subtle changes. These computational predictions were directly tested in a subsequent behavioral experiment.

### Modulation of PV^+^ interneurons alters memory interference

To experimentally validate the model predictions, we designed a new behavioral protocol that we called multi-arena NOL task. This task was specifically designed to assess how manipulations of PV^+^ inhibitory activity affect memory interference in conditions of high memory load. The multi-arena NOL task evaluates an animal’s ability to maintain multiple distinct memories through four unique contexts, each defined by multiple environmental cues (color, shape and landmarks), different objects and specific object-context-location associations (Fig. 4a). The complete experimental protocol was conducted over three consecutive days. Analysis of the averaged DI revealed significant group differences (Fig. 4b). Notably, PV-Gi animals – where inhibition was reduced and discrimination enhanced in low memory-load experiments (Fig. 3b, g) – showed in this task a pronounced impairment compared to controls, whereas PV-Gq animals, with enhanced inhibition, exhibited intermediate performance. These behavioral results mirror the model’s predictions: insufficient inhibition increases memory interference, while high inhibition supports more stable memory traces at the cost of discriminability, thereby impacting animals’ behavior.

**Figure 4.**
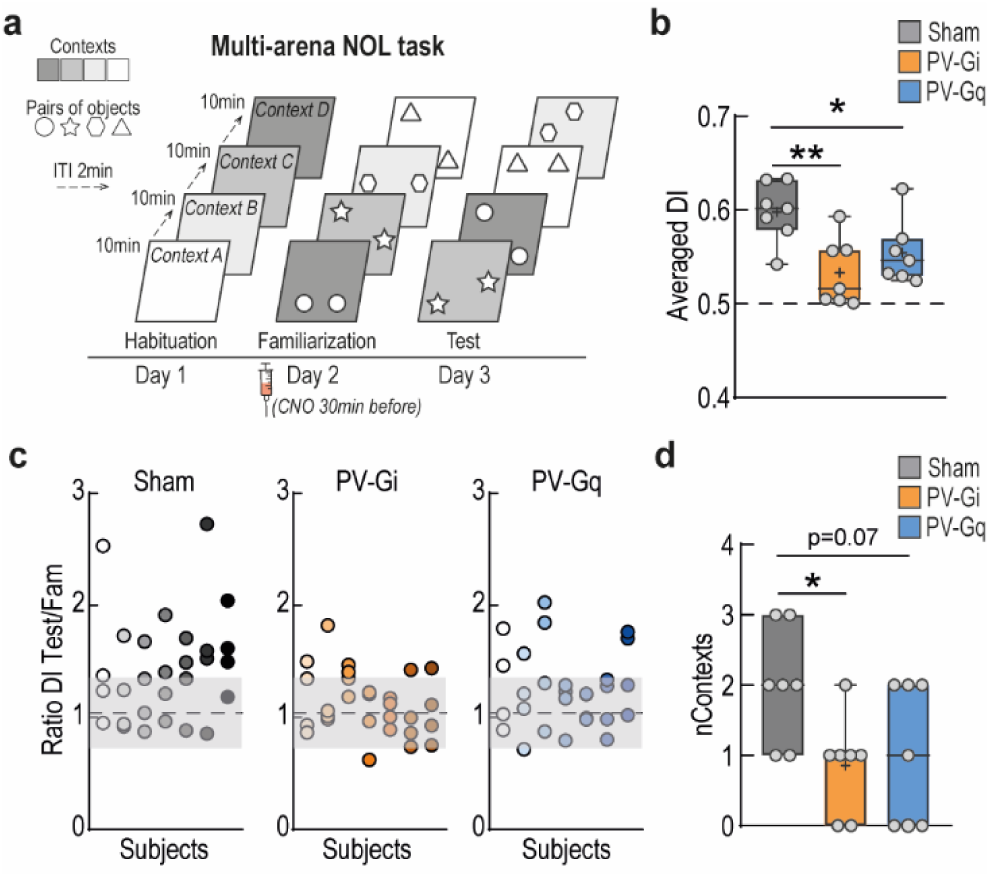
Impact of DG-PV modulation on memory discrimination under high memory load conditions. **(a)** Experimental protocol for multi-arena NOL task, describing the three-day task designed to assess memory encoding in higher memory-load conditions, by introducing 4 different arenas. **(b)** Comparison between groups: averaged DI (aDI) is calculated from the mean performance across all four contexts for each subject for Sham (n = 7), PV-Gi (n = 7), and PV-Gq (n = 7) groups. **(c)** DI ratio between test and encoding sessions in each context obtained for each animal and sorted by group. Subjects are displayed along the X-axis, and the four contexts explored per subject are represented in vertical columns (symbols filled with matching color tones). Shaded area represents the mean ± 2*STD from the non-learning baseline obtained from PV-Gq animals in the NOL task. **(d)** Estimated number of contexts learned by each group of animals, defined as the number of contexts in which the DI ratio exceeded a reference threshold. This threshold was set as the mean plus two standard deviations of DI ratios from Gq animals in the NOL task (see Materials and Methods). All statistical values are detailed in Table S1. * = p<0.05, ** = p<0.01.

We further estimated the number of contexts successfully encoded by each animal. This was done by comparing individual DI ratios for each context against a threshold derived from variability in non-learning conditions (Fig 4c; see Material and Methods). The number of contexts learned varied across groups, with a significant reduction in PV-Gi animals compared to Sham controls, and a trend toward reduction, but at intermediate levels, in PV-Gq animals (Fig 4d).

Together, these findings support the idea that, as memory load increases, maintaining PV^+^-mediated inhibition within a functional range is critical for preserving the fidelity of episodic memory representations. Departure from this balance – either through excessive disinhibition or hyperinhibition – compromises the system’s ability to store and retrieve multiple distinct memories.

Overall, the convergence of computational and experimental results suggests that PV⁺ interneurons in the DG regulate the position of the system along a sensitivity–consistency continuum of memory representation.

## Discussion

By manipulating PV^+^ cells activity in the DG, we show that reduced perisomatic inhibition during spatial memory encoding enhances discrimination. This effect is consistent with simulation results, which suggest that heightened sensitivity to subtle input differences can promote discrimination by generating distinct activity patterns. Reduced inhibition would increase the chance that small input variations activate different neuronal populations. As a result, similar inputs generate more distinguishable representations. Importantly, the same computational model predicted a downside: increased susceptibility to synaptic interference and, therefore, reduced memory stability. To test this, we developed the multi-arena NOL task, designed to challenge memory consistency across multiple contexts. Animals with reduced inhibition during encoding were selectively impaired in this task, supporting the idea that enhanced sensitivity may compromise resistance to interference.

Conversely, sustained activation of PV^+^ cells during memory encoding impaired discrimination performance later during recall. Both behavioral and computational modeling suggest that this effect is not due to a failure of memory formation per se, but rather to diminished pattern discrimination: strong inhibition limits the network’s ability to form distinct representations when input differences are subtle, leading to increased generalization and preservation of existing memory traces. In line with this, animals with increased PV^+^ activation during encoding in the multi-arena NOL task were still able to learn some spatial contexts, likely those in which object displacements were more salient. These findings suggest that while excessive inhibition may impair the ability to encode precise spatial differences, it does not fully prevent memory formation when discriminative demands are lower.

Together, our study underscores a fundamental computational trade-off in the DG: reduced inhibition enhances discrimination by increasing the system’s sensitivity to inputs with high similarity, but heightened sensitivity comes at the expense of stability, rendering representations more vulnerable to interference. In contrast, stronger inhibition promotes consistency and resistance to interference but limits the system’s ability to update the memory base with fine-grained discriminative differences. Such dynamic tuning of inhibitory strength may reflect an adaptive mechanism by which the hippocampus shifts between sensitivity- and consistency-oriented modes depending on task demands or cognitive state. Importantly, alterations in the excitation/inhibition balance have been linked to a range of neuropsychiatric and neurodegenerative disorders, including schizophrenia, depression and age-related cognitive decline ^20-22^. Therefore, our findings not only reveal a circuit-level mechanism that modulates the trade-off between memory sensitivity and consistency but also offer insight into how disrupted inhibition control may contribute to dysfunction.

Episodic memories are initially acquired within the hippocampus–entorhinal cortex network through rapid synaptic plasticity and are subsequently consolidated in neocortical regions for long-term storage ^5, 23-29^. The interaction between the hippocampus and the prefrontal cortex plays a crucial role during the encoding of new memories and is essential for their later recall ^30^. Modulation of hippocampal inhibitory circuits, as performed in our study—particularly during the encoding phase—is therefore expected to influence this broader memory network. Indeed, we have previously shown, using brain-wide imaging ^31^, that activity modulation in the dentate gyrus ^32-34^ and CA3 ^35^ regulates functional connectivity across a wide network of memory-related structures, including the hippocampal formation, the prefrontal cortex, and the nucleus accumbens.

Our computational model was designed to capture essential principles of how inhibition shapes memory computations within a cortico-hippocampal system, specifically the trade-off between discrimination and stability in the context of novelty-driven decision making. Rather than aiming for detailed biological realism, we prioritized conceptual clarity and generalizability. This level of abstraction allowed us to isolate the effects of inhibitory tone on representational dynamics and memory-guided behavior without relying on complex parameter tuning or intricate circuit assumptions that could reduce interpretability. While more detailed models, such as biophysically realistic spiking networks, can capture finer aspects of memory and decision making ^36-38^, they often involve fitting numerous parameters, which can limit their explanatory power and increase the risk of overfitting ^39^. By contrast, our model balances between simplicity and explanatory depth. Conceptual models of this kind have been widely and successfully used to uncover the mechanisms behind memory and decision making ^40-43^. Our model provides a minimal yet insightful computational framework for understanding how inhibitory modulation may influence memory-based computations, offering a plausible mechanistic perspective on systems-level memory function.

The multi-arena NOL task also warrants consideration. While it enables assessment of memory consistency across multiple contexts, it does not disentangle the temporal dynamics of interference. Specifically, it remains unclear whether interference arises from the weakening of earlier-encoded memories or from a reduced capacity to store new ones once others are consolidated. This limitation is not specific to our design but reflects a broader challenge in behavioral paradigms aimed at probing how sequential experiences compete for representation ^44, 45^. In our task, animals explore a sequence of contexts and are later tested in a shuffled configuration, introducing unavoidable interference between encoding and recall. Thus, although our findings clearly indicate that reduced inhibition impairs memory stability across contexts, the current design cannot determine whether early or late memories are more affected. Resolving this question will require dedicated experiments and novel behavioral strategies that isolate specific memory phases without repeated exposure to the same episodes.

Overall, flexibility and stability are both essential for adaptive behavior: flexibility allows the memory system to incorporate new information and adjust to changing environments, while stability preserves existing knowledge against interference ^46^. Striking the right balance between these demands is critical. Excessive flexibility can lead to volatility, with memories overwritten by each novel event, whereas excessive stability can result in rigidity, preventing the learning of important new information. Such rigidity has been hypothesized to underlie the persistent cognitive inflexibility observed in patients with several psychiatric conditions ^47, 48 49, 50^. Here, by linking PV^+^ interneuron modulation to the sensitivity– consistency trade-off, we provide a plausible circuit mechanism by which the brain navigates the continuum between flexibility and stability during learning.

## Methods

### Animals

All animal experiments were approved by the Animal Care and Use Committee of the Instituto de Neurociencias de Alicante (Alicante, Spain) and complied with the Spanish law (53/2013) and European regulations (EU directive 2010/63/EU).

All animals were two months old at the beginning of the experiments and were randomly assigned to different experimental groups (see below). We excluded animals showing low exploration behavior (see NOL task). No significant differences between sexes were found and thus the data were pooled. Mice were bred in house from the line 129P2-Pvalbtm1(cre)Arbr/J (Jackson Laboratories, RRID: IMSR_JAX:008069) and housed in groups (3-5), with 12-12 h light/dark cycle, lights on at 8:00, at room temperature (23 ± 2 °C) and free access to food and water.

### Viral injections

#### Surgery

Mice were anaesthetized with isoflurane (Laboratorios Esteve, Murcia, Spain) 4%, 0.8L/min oxygen for induction, 1-2%, 0.8L/min oxygen for maintenance, and then fixed in a stereotaxic device (David Kopf Instruments, California, USA) over a heating pad at 37°C. Coordinates for targeting the injections in the hilus of the DG, from Bregma, were -2mm AP, ± 1.4 mm LM, +2mm DV ^51^. After opening the skin, we opened a 700μm Ø trepan with a milling cutter (Ref: FST 19007-07, Finest Science Tools, FST, Heidelberg, Germany) attached to a cordless micro drill (Ref: 58610V, Stoelting Co., Illinois, USA). Then we gently introduced a micropipette (Ref: 4878, World Precision Instruments, WPI, London, UK) through the trepan, into the brain. Micropipette was filled with mineral oil and viral vectors containing Designer Receptors Exclusively Activated by Designer Drugs (DREADDs). Oil and viral vectors were separated with c.a. 1 μl of air. Micropipette was attached to a pump infusion Nanoliter 2010 Injector (WPI) coupled to the stereotaxic frame and was previously pulled in a P-2000 puller (Sutter Instruments Company, California, USA). We placed the tip of the micropipette in the hilus and very slowly injected 0.5 μl of viral vectors per hemisphere. After injection was completed, we waited 10 min before retracting the micropipette 200 μm, then waited another 10 min and finally removed gently and completely the pipette. After closing the skin with silk thread we administered subcutaneously analgesic, buprenorphine (Dechra Veterinaria Products SLU, Barcelona, Spain), and antibiotic, Syvaquinol (Laboratorios Syva S.A.U., León, Spain), to the mice and we controlled their recovery, injecting additional doses of analgesic if necessary.

#### DREADDs, Designer Receptors Exclusively Activated by Designer Drugs

To modulate PV interneuron activity in the PV-cre mice we used DREADDs (UNC Gene Therapy Vector Core, University of North Caroline, North Caroline, USA; Viral Vector Facility, Neuroscience Center Zurich, Switzerland). To inhibit PV cells we injected adeno-associated virus 5 (AAV5)-human synapsin1 (hSyn)-DIO-hM4D(Gi)-mCherry, and for activation we used the AAV5-hSyn-DIO-hM3D(Gq)-mCherry ^17^. Moreover, for c-Fos expression we also used the control virus AAV5-hSyn-DIO-mCherry. The specificity of the expression in PV^+^ cells was confirmed immunohistologically (Fig. S1). After viral injection we allowed 3-4 weeks for DREADDs expression. Clozapine-N-Oxide (CNO, 1mg/kg, i.p in saline 0.9%; ENZO Life Science Inc., New York, USA), was used for DREADDs activation ^17^.

### Electrophysiology

All electrophysiological protocols were recorded in the same animals before and after CNO injection (n=6 PV-Gi, n=6 PV-Gq and n=6 Sham). Mice were anaesthetized with urethane (1.4 g/kg, i.p.) and recording and stimulating electrodes were implanted following stereotaxic standard procedures. A bipolar stimulating electrode (WPI, ref. TM53CCNON) was placed in the perforant pathway with the following coordinates relative to bregma: -4.3 mm anteroposterior (AP), +2.5 mm mediolateral (ML), +1.4 mm dorsoventral [DV], and at an angle of -12°. The DV position was adjusted individually for each animal to optimize the evoked response. The recording probe had a single shank and 32 channels (50 μm contact spacing; NeuroNexus Technologies, Michigan, USA). It targeted the hippocampus (-2 AP, +1.5 ML, -2 DV), including CA1 and DG regions.

Electrophysiological recordings were made before and 90 min after CNO injection (1 mg/kg, i.p.). After post-CNO recording, we induced LTP using a standard theta burst protocol: six 400 Hz-trains of pulses delivered at a 200 ms interval, repeated six times with 20 s inter-train interval (Stäubli et al., 1999). Evoked potentials were recorded again 60 min after LTP induction. We applied a stimulation intensity adjusted to 80% of the maximum PS.

Signals were filtered (0.1-3k Hz), amplified and digitized (20-32 kHz acquisition rate), with data analyzed offline using Spike2 software (Cambridge Electronic, Cambridge, UK) or MatLab (The MathWorks Inc., Natick, MA, USA).

To quantify the electrophysiological responses, we measured the EPSP as the maximum slope of the evoked potential in the molecular layer (for DG). The PS was measured as the amplitude of the fast negative potential recorded in the hilus of the DG. In the PV-Gi animals, in which CNO administration caused a large increase in PS amplitude, the EPSP was measured more distal from the granule cell soma layer (in the outer third of the molecular layer) to minimize the volume-conducted artefact of the PS, enduring more accurate EPSP slope measurements.

### c-Fos experiments

To quantify the impact of PV modulation on c-Fos expression after encoding, 22 animals (8 PV-Sham, 8 PV-Gi, 6 PV-Gq) were allowed to freely explore a novel environment (a 50x50x30 cm arena containing spatial cues and two identical objects) and sacrificed 60 min later. Perfusion of the animals and tissue slicing were performed as described above. Eight to ten serial slices, covering the hippocampus, PFC and NAc were processed for c-Fos immunostaining (primary antibody: guinea pig anti-cFos, 1:1000, ref. 226004, Synaptic Systems, Göttinger, Germany; secondary antibody: Alexa 488 donkey anti-guinea pig, 1:500, ref. 706-545-148, Jackson ImmunoResearch, Suffolk, United Kingdom). The slices were mounted and used in subsequent quantitative analysis.

Quantitative analysis of c-Fos positive nuclei was performed offline using 12-bit grayscale images acquired with a Leica DM4000 fluorescence microscope at 10x/0.25 dry objective. Images were analyzed using a Neurolucida software (MBF Bioscience, Williston, VT USA). The ROIs were manually delineated following the Franklin and Paxinos mouse brain atlas ^51^. Analysis was conducted using Icy Software ^52^ in a semi-automatic manner. The detection threshold for positive nuclei was set for each brain region, considering the average nuclei size and a signal-to-noise ratio greater than 23%, in accordance with the Rayleigh criterion for resolution and discrimination between two points. Animals with misplaced viral infections were excluded from all the analysis.

### Histology

Animals were perfused with 50 ml of 37°C saline and 50 ml of 4% paraformaldehyde (PFA; BDH prolabo, VWR chemicals, Lovaina, Belgium). Brains were extracted and stored in 4% PFA at 4°C for at least 24h. After fixation, brains were sectioned using a vibratome (VT 1000S, Leica, Wetzlar, Germany) into 50 μm slices. Validation of anatomical coordinates for recording and stimulating electrodes was done in DAPI stained slices. Viral infection efficiency and specificity were quantified in 6-8 hippocampal containing slices following standard immunohistochemistry protocols. For PV labeling, primary antibody (mouse anti-PV, 1:2000, ref. 235, Swant, Switzerland) and secondary antibody (Alexa 488 goat anti-mouse, 1:500, ref. A11029, Life Technologies; USA) were applied. Virus expression was further visualized using the mCherry reporter co-labelling.

### Behavior

To assess the impact of PV modulation in spatial memory, we used a series of behavioral tasks. To quantify the animals’ exploratory behavior, we used a custom MATLAB-based code developed in the lab, which allowed animal tracking and automatic quantification of exploratory behavior.

#### Novel Object Location (NOL) task

To assess spatial memory, a total of 121 mice performed the NOL task. After a period of handling, mice were introduced into an empty arena (50x50x30 cm) with spatial cues and soft illumination (luxes: 23 in the center ± 2 in the corners). They were allowed to freely explore the arena for 2 periods of 5 min (habituation phase). Twenty-four hours later, two identical objects were introduced in opposite corners of the arena (13.5 cm from the walls), and the animals were allowed to explore them (familiarization or encoding phase). Familiarization ended when an animal accumulated 20 s of exploration of both objects or after 10 min in the arena ^19^. Animals that explored the objects less than 8 s during the 10 min of the familiarization phase were excluded from the study (*a priori* exclusion criteria). Another 24 hours later, the mice were reintroduced in the same arena, but one of the objects was displaced to a new location (another corner, 13.5 cm from the walls and 15 cm from the other object). Mice were again allowed to explore under the same criteria (test or retrieval phase) (Fig. 2a).

To assess spatial memory, we calculated a DI, measured as the proportion of time spent exploring the displaced object relative to the total exploration time of both objects. Using this behavioral protocol, we administered CNO or its vehicle at different time points to modulate PV interneuron activity: during encoding (CNO or saline injection 90 min before the familiarization phase; n=38), during consolidation (CNO injection 10 min after familiarization; n=42) or during retrieval (CNO injection 90 min before the test phase; n=41) (Fig. 2a). For encoding and control experiments, the same animals were used. Different groups were used for consolidation and retrieval experiments.

In addition to spatial memory, we measured and quantified locomotor parameters such as distance traveled (Fig. S2).

#### Sequential NOL task

To assess spatial memory discrimination, we implemented an extension of the NOL task. A total of 8 PV-Gi mice underwent identical habituation and encoding protocols as in the NOL task. During the test phase, animals were reintroduced to the arena where one object was displaced 10 cm (rather than the standard 15 cm). Following a 1-minute interval, the same object was displaced an additional 5 cm. All animals were tested under both experimental conditions (CNO vs vehicle) using a within-subjects design, with treatments administered 90 min prior to familiarization. Conditions were separated by a 1-week interval to ensure complete drug clearance and memory washout.

#### Multi-Arena NOL Task

To assess spatial memory capacity, 21 animals (n=7 PV-Gi, n=7 PV-Gq and n=7 Sham; see Suppl. Exp. Proc.) performed the multi-arena novel object location task (multi-arena NOL) task (Fig. 4a), designed as an extension of the NOL task. This task evaluated the ability of mice to encode and recall object locations across multiple contexts.

Mice were sequentially exposed to four distinct arenas, each defined by unique combinations of shape, texture and material, to provide distinct contextual cues. In each arena, they encountered a different pair of identical objects, varying in shape, size, material and texture to strengthen object-context associations. Object positions were arranged to differ maximally between contexts across encoding and retrieval sessions. The order of arena presentation varied across sessions to prevent sequence learning mechanisms or strategies.

The behavioral apparatus consisted of a rectangular corridor (20 cm length x 12 cm height x 10 cm width, made of white methacrylate), connecting two distinct contexts, one at each end. Entrance gates were manually opened by the experimenter to allow the mice free access to one context at a time.

The behavioral protocol followed the same 3-day time flow followed in the NOL task: habituation,familiarization and test sessions. Each session consisted of four trials (one visit per context). On Day 1 (habituation), mice sequentially explored all four contexts (*A→B→C→D*) for 10 min each, with 2-min waiting periods in the central corridor before the session and between contexts. On Day 2 (encoding), mice received injection of CNO (1 mg/Kg, i.p.) 60 min before the session. They explored the contexts in a different order than during habituation (*D→B→C→A*), and encountered two identical objects at fixed locations within each arena. On Day 3 (retrieval), one object in each context was displaced to a new location, and the order of context exposure was altered again (*C→D→A→B*). The other object remained stationary. Each session lasted 50 min per animal (4 trials x 10 min + waiting periods).

Spatial memory capacity was assessed by measuring the number of contexts in which the mice preferentially explored the displaced object over the stationary one, indicating their ability to discriminate object displacement. In addition, we used an averaged DI (*aDI*), which takes into account the mean preference of the animal for all contexts:

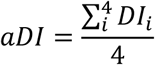

#### Calculation of DI range for non-learning condition

To estimate the number of contexts learned by each group, we calculated the DI ratio between test and encoding for each context. As a reference for non-learning, we used the distribution of DI ratios from Gq animals in the NOL task – known to perform poorly - and defined the upper bound of this reference range as the mean plus twice the standard deviation. DI ratios exceeding this upper bound were considered indicative of learning. For each animal, we counted the number of contexts in which the DI ratio surpassed this reference, providing an estimate of how many contexts had been successfully learned.

### Computational model

The computational model consisted of two modules ^43^: one module with functionalities related to the hippocampus, such as learning and pattern separation, and another module with functionalities related to the prefrontal cortex, such as decision making.

The hippocampal-related module was modeled as a neuronal layer with convergent projection from a “sensory” representation onto a memory space, with local competition between neurons ^42, 53^. Local competition was implemented using a k%-max winner-take-all mechanism ^9^, which simulates different levels of inhibition. In this mechanism, only the neurons whose activity falls within a k% of the maximum neuronal activity are activated. Higher values of k% reduce inhibition, increasing the probability of a neuron being active, while lower values increase inhibition.

The activity of 200 neurons simulates the “sensory” input pattern projected to the memory space. Each neuron is assigned a random activity value, drawn from a uniform distribution and normalized so that the total sum is 1. These neurons connect with a probability of 25% to 500 neurons in the memory space. During learning, the synaptic weight between an active presynaptic neuron (*a*_*i*_) and an active postsynaptic neuron (*a*_*j*_) increases based on the activation values of both neurons and the learning rate (*L*_*r*_):

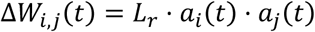

After each training step, the weights are normalized to their mean value. In the simulations, we used a learning rate of *L_r_* = 0.05 to produce stable representations of learned patterns ^43^. A postsynaptic neuron (*a*_*j*_) in the memory space becomes active only if its input exceeds k% of the maximum input activity across all neurons in the layer. If this is the case, the neuron’s activity is set as the difference between its input activity and the k% from the maximum input activity, normalized by the maximum activity of all active neurons.

The hippocampal-related module was selected based on its previously proven validity to explain some of the dynamics taking place in the hippocampus, such as local competition or rate remapping ^9, 42^. The sum of the hippocampal neurons’ activity is projected onto the module associated with prefrontal cortex functionalities, where the decision is made.

The decision-making module is a mean-field approximation of a realistic network of integrate-and-fire neurons ^54^. The rate-based model is a well-established approximation that allows the study of neuronal activity while reducing the computational expenses of modeling individual neurons. Moreover, it is supported by the fact that neurons are organized in neuronal populations with similar properties ^55^. The model consists of two groups of excitatory neurons with recurrent connections (“explore object” and “explore arena” groups) that compete through mutual inhibition ^56, 57^. The activity of the excitatory groups follows the dynamics:

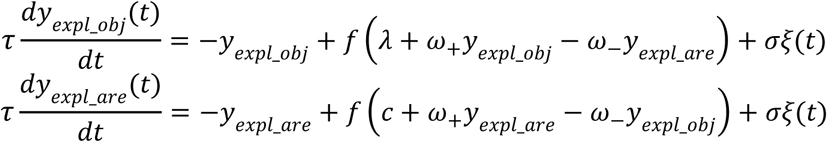

where *y* represents the activity of each group, *ω* is the weight of the connections (*ω*+ recurrent; *ω*− inhibitory), *λ* represents the input from the hippocampal-related module to group “explore object”, c represents a constant input to the “explore arena” group, which mimics the activity from a learned (familiar) input pattern, *σξ* represents the network fluctuations and *f(.)* is a sigmoidal function of the form:

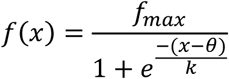

where *f*_max_ defines the maximum activity, i.e. the activity at which the neuronal network saturates. The network fluctuations are simulated through a Gaussian noise with mean 0 and variance 1 (*ξ*), which simulates having a finite set of neurons, and a scale factor *σ* = 0.01 spikes·s^-1^. In the simulations, we used: *τ* = 20 ms, *ω*_+_1, *ω*_−_= 1, *f_max_* = 40 spikes⋅s^-1^, *θ* = 4 spikes⋅s^-1^, and *k* =11 spikes⋅s-1. Within the prefrontal-related module, the strength of the connections was kept constant during the simulations.

To simulate the DI during the testing sessions, the input to the “explore arena” group in the decision-making module was set constant, simulating the output of the hippocampal module for familiar patterns. The “explore arena” group received the output of the hippocampal module as the sum of the activity of its neurons. The input activities were scaled by a factor of 0.1. A decision towards exploring the object was made when the “explore object” group won the competition. The DI was calculated as the ratio between the number of decisions towards exploring the object when the input pattern was novel (not learned) and the number of decisions made towards exploring the object when the pattern was familiar (learned) or novel (total cases).

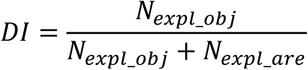

In the simulations, we used 20 “sensory” input patterns to train the hippocampal-related module, except when investigating memory stability, where the number of patterns varied. During learning, each pattern was presented as input for 50 trials and was subsequently referred to as a familiar pattern. After learning, the same or a specifically modified familiar pattern was presented for 30 runs at the input to test the model. These modified patterns were generated by introducing different levels of noise to the original input pattern, resulting in patterns with different degrees of similarity to the familiar one. Similarity between sensory representations was quantified as the correlation between the activity vector of familiar and modified input patterns. For each level of input similarity and inhibition (competition) levels, the entire procedure was repeated 10 times. The similarity of memory representations was then quantified as the correlation between the output evoked by a familiar pattern and the outputs evoked by gradually modified versions of that pattern.

To simulate the sequential NOL task, the model first learned 20 input patterns over 50 runs, as previously. The model was then tested with these learned familiar patterns (familiar condition) and with noisy variants exhibiting a correlation of 0.8 with one of the familiar patterns. These noisy patterns were subsequently used for a second learning phase (10 runs), after which the model was tested again using newly generated noisy patterns with a correlation of 0.6 between familiar and those new patterns.

To evaluate the model’s robustness to noise, we varied the strength of the connections between the input and memory space within the hippocampal-related module. Each connection weight was modified by adding a pseudorandom value between 0 and 1, scaled by a factor ranging from 0 to 5 in steps of 1.

Memory interference was assessed by increasing the number of patterns from 50 to 1000, in increments of 50, for each level of competition.

### Statistical analysis

The statistical analysis was done using GraphPad Prism 7 software (GraphPad Software Inc., La Jolla, CA, USA) or SPSS v20 software (IBM, New York, USA). After an exploratory analysis for the presence of outlier values, we checked the skewness and kurtosis of the data before testing their statistical distribution. Parametric-test requirements, including normality (D’Agostino-Pearson test and Shapiro-Wilk test) and homoscedasticity (F of Levene test) were tested. All the data fulfilled parametric criteria, unless otherwise specified. In most analyses we applied two-way ANOVA, with a *group* factor with 3 levels (Sham, PV-Gi and PV-Gq), and a *time* factor with 2 levels (before and after CNO injection). In case of c-Fos analysis, we applied one-way ANOVA with group as a factor with the same 3 levels as above. We applied a Sidak *post hoc* analysis for multiple comparisons with adjusted alpha. Effect size was calculated with partial *eta* square (ηp^2^ as SSeffect / (SSeffect +SSerror), and the power of the effect (1-β) using GPower software (University of Düsseldorf, Düsseldorf, Germany). Effect size is ^2^considered large if ηp^2^ > 0.13. If, by any reason, partial *eta* square could not be used, then we express ^2^r indicator, being similar to *eta*, but requiring > 0.30 values to indicate large effects. The power of the effect (1-β) is taken as an invert indicator of committing a type II error. The results of the statistical analysis are detailed in Supp. Tables numbered according to the number of the Figures to which they correspond.

Experimental data were plotted using GraphPad Prism 7 software (GraphPad Software Inc.) and figures were composed using Adobe Illustrator software (Adobe Systems Incorporated, San José, California, USA).

## Acknowledgements

We are grateful to Professor Richard G.M. Morris for his insightful comments on the manuscript. We also thank Alejandro Sempere and César Rennó-Costa for discussions and feedback at different stages of the work. The excellent technical assistance by Analía Rico Rodríguez is greatly acknowledged. This research was financially supported by the Spanish Ministerio de Ciencia e Innovación, Agencia Estatal de Investigación Grant PID2021-128158NB-C21, funded by MICIU/AEI/10.13039/501100011033 and ERDF/EU, awarded to SC; by Grant RYC2021-035061-I and Grant PID2022-141173NA-I00, funded by MICIU/AEI/10.13039/501100011033 and by the European Union NextGenerationEU/PRTR and ERDF/EU, respectively, awarded to EM. We also received support from the Excellence Grant CIPROM/2022/15 to SC and EM, funded by the Generalitat Valenciana and Next Generation EU. We also acknowledge the Program for Centres of Excellence in R&D Severo Ochoa, Agencia Estatal de Investigación (CEX2021-001165-S) funded by the MICIU/AEI/10.13039/501100011033.

## Supplemental Information

Supplemental Experimental Procedures

### Behavior

We used a series of experimental tasks to assess the influence of PV^+^ cell modulation on non-spatial memory and anxiety levels.

#### Elevated Plus Maze (EPM)

To assess potential anxiety-related changes induced by PV-cell activity modulation in the DG, we conducted the elevated plus maze (EPM) test. One week after the NOL task, animals were injected either with CNO or saline, and placed in the EPM 90 min later. The maze consisted of two open and two closed arms (Fig. S3D). Behavior was video-recorded and tracked using commercial software (Smart Video Tracking Software, Panlab, Barcelona, Spain). Anxiety levels were evaluated by comparing the time spent in the open *vs.* closed arms. Additional parameters, such as number of arm entries and distance traveled, were also analyzed (Fig. S2E).

Following behavioral evaluation, mice were assigned to groups for further electrophysiological or fMRI experiments.

**Figure S1.**
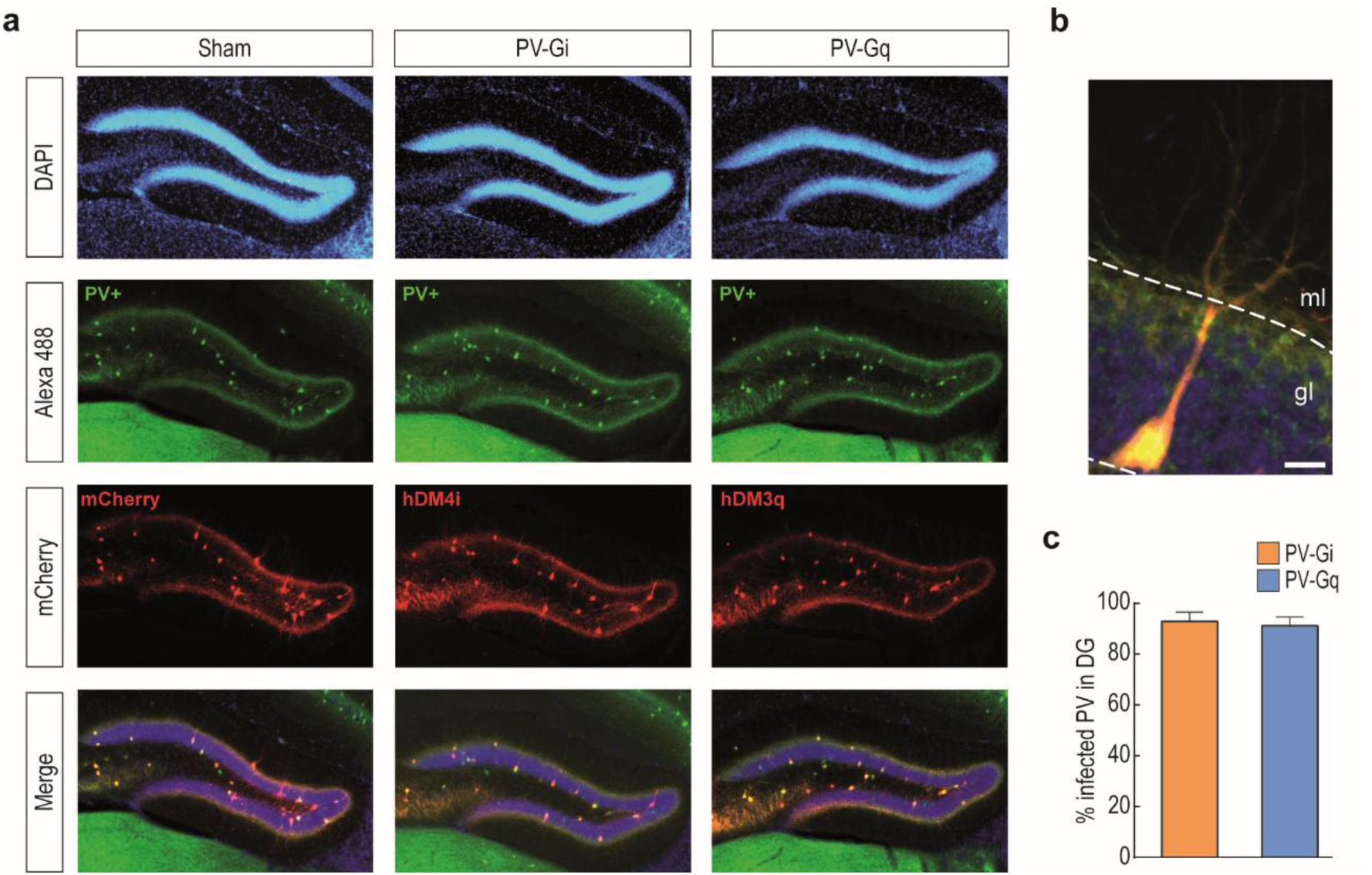
Expression of DREADDs in the DG of PV-cre animals. **(A)** Representative pictures showing the reporter of the virus infection (mCherry, red), immunolabelled against PV protein (green) and counterstained with DAPI (cell nuclei, blue). Scale bar: 100 μm. **(B)** Zoom in of an infected PV^+^ cell (white square in A). Scale bar: 10 μm. **(C)** Efficiency of DREADDs expression in the DG expressed as the % of infected PV-cells in PV-Gi (orange) and PV-Gq (blue) animals. Data show mean ± SEM.

**Figure S2.**
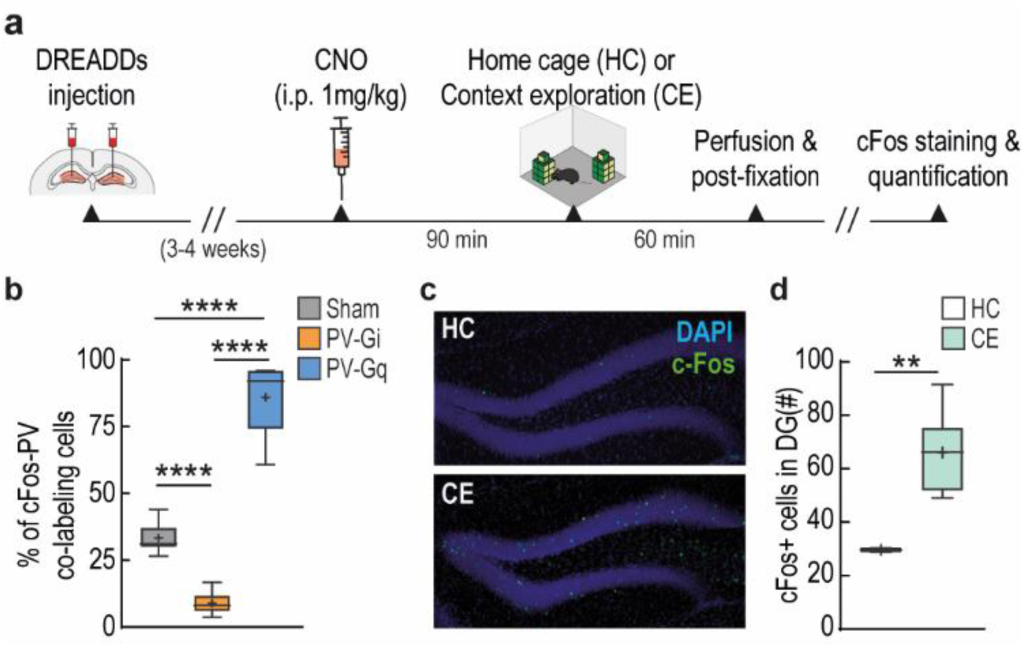
Quantification of activated c-Fos+ cells. **(a)** Temporal sequence of the experimental protocol for c-Fos+ cells quantification. After 90 min post-CNO injection, animals were either allowed to explore a novel arena with 2 objects (encoding phase of a NOL task) or maintained in their home cage. **(b)** Quantification of modulated PV+ cell activity, expressed as the percentage of PV+ cells in the DG that are double stained for c-Fos. **(c)** Example of activated c-Fos+ cells in DG in home cage (HC) and after NOL encoding phase (context exploration, CE). Scale bar: 100 μm **(d)** Number of activated c-Fos+ cells in the DG during the NOL encoding phase vs. HC. Scale bar: 100 μm. All statistical values are detailed in Table S2. **p<0.01, ****≤0.0001.

**Figure S3.**
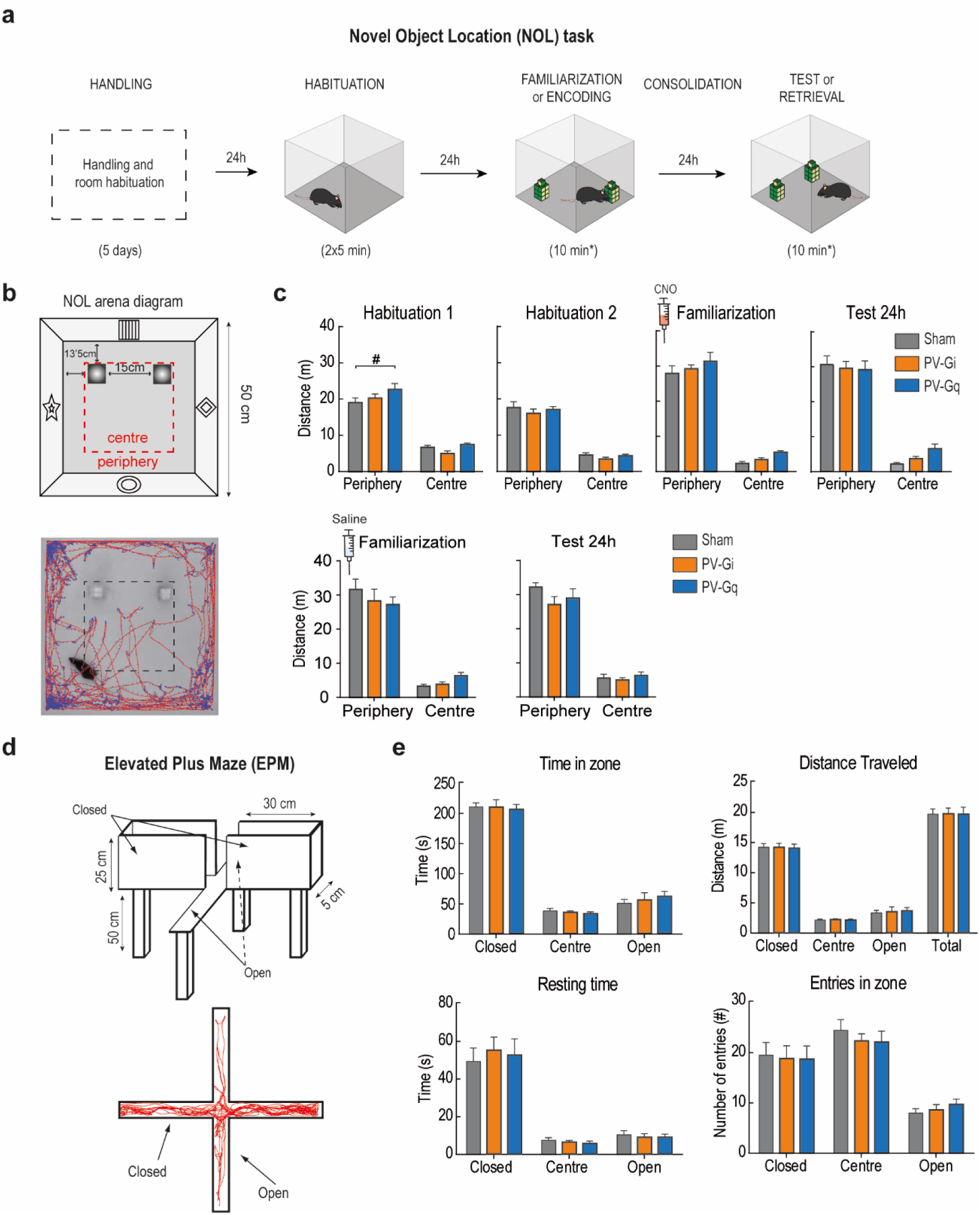
Motor activity and anxiety. **(A)** Diagram of the Novel Object Location (NOL) protocol followed. **(B)** Detailed diagram of the NOL arena. Example of one mouse tracking within a test session (bottom). **(C)** For evaluating locomotor activity, with and without PV modulation, the travelled distance in each session was analyzed for the three groups of mice. **(D),** Diagram of the Elevated Plus Maze used for evaluating anxiety (top). Example of one trial (bottom). **(E)** Anxiety levels as measured by time spent, distance travelled, resting time and entries per zone. All statistical values are detailed in Supp. Table S2.

**Table S1.**
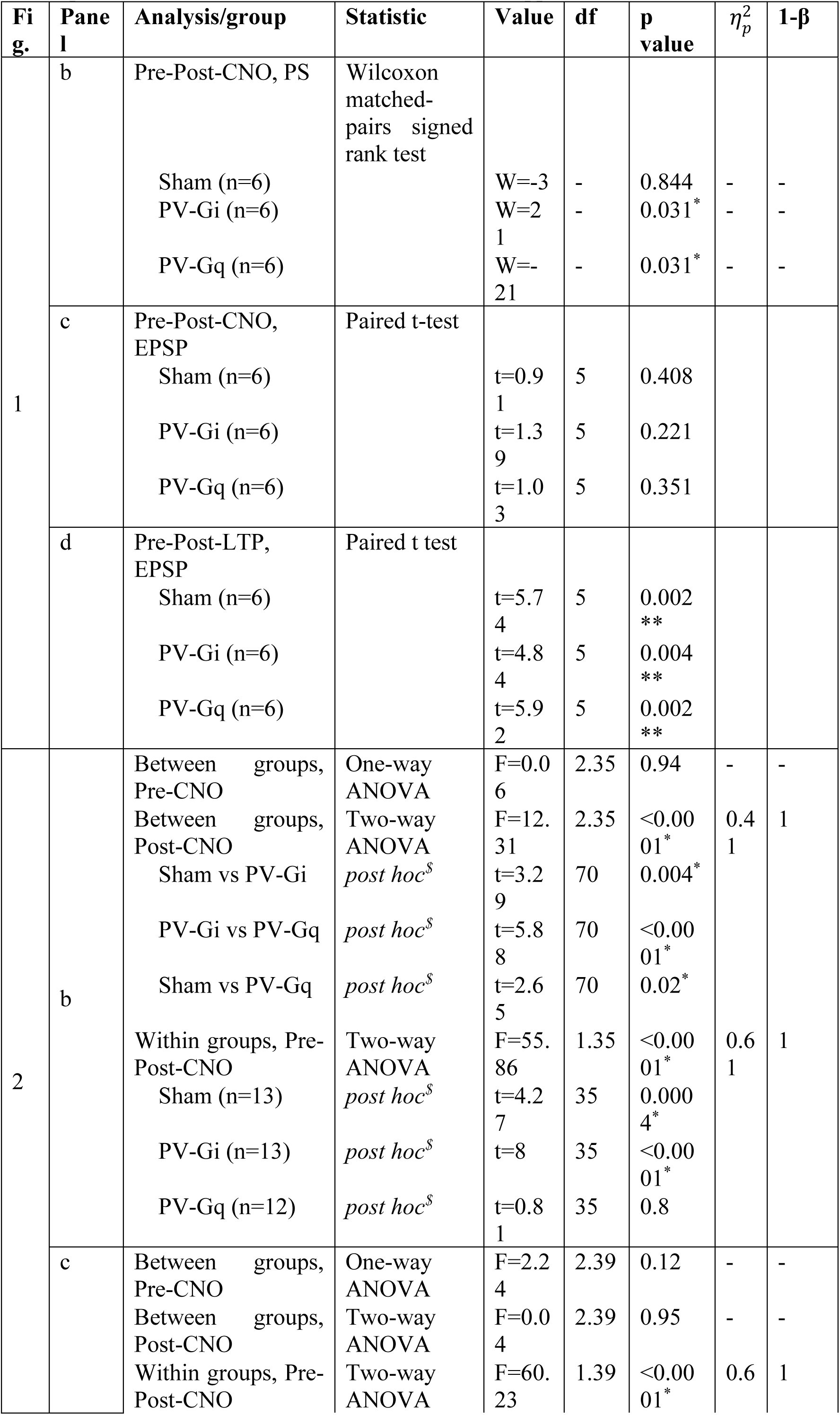
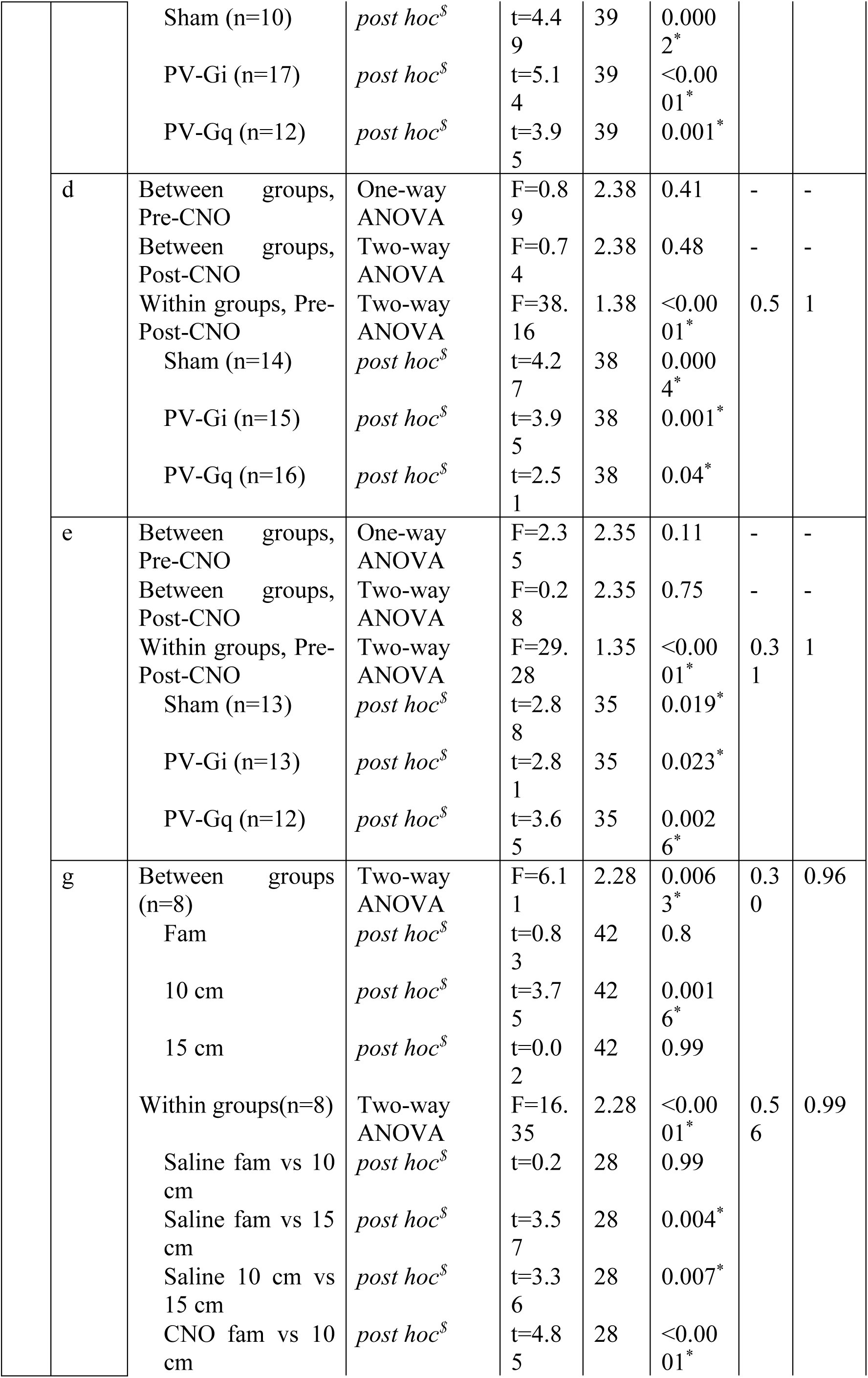
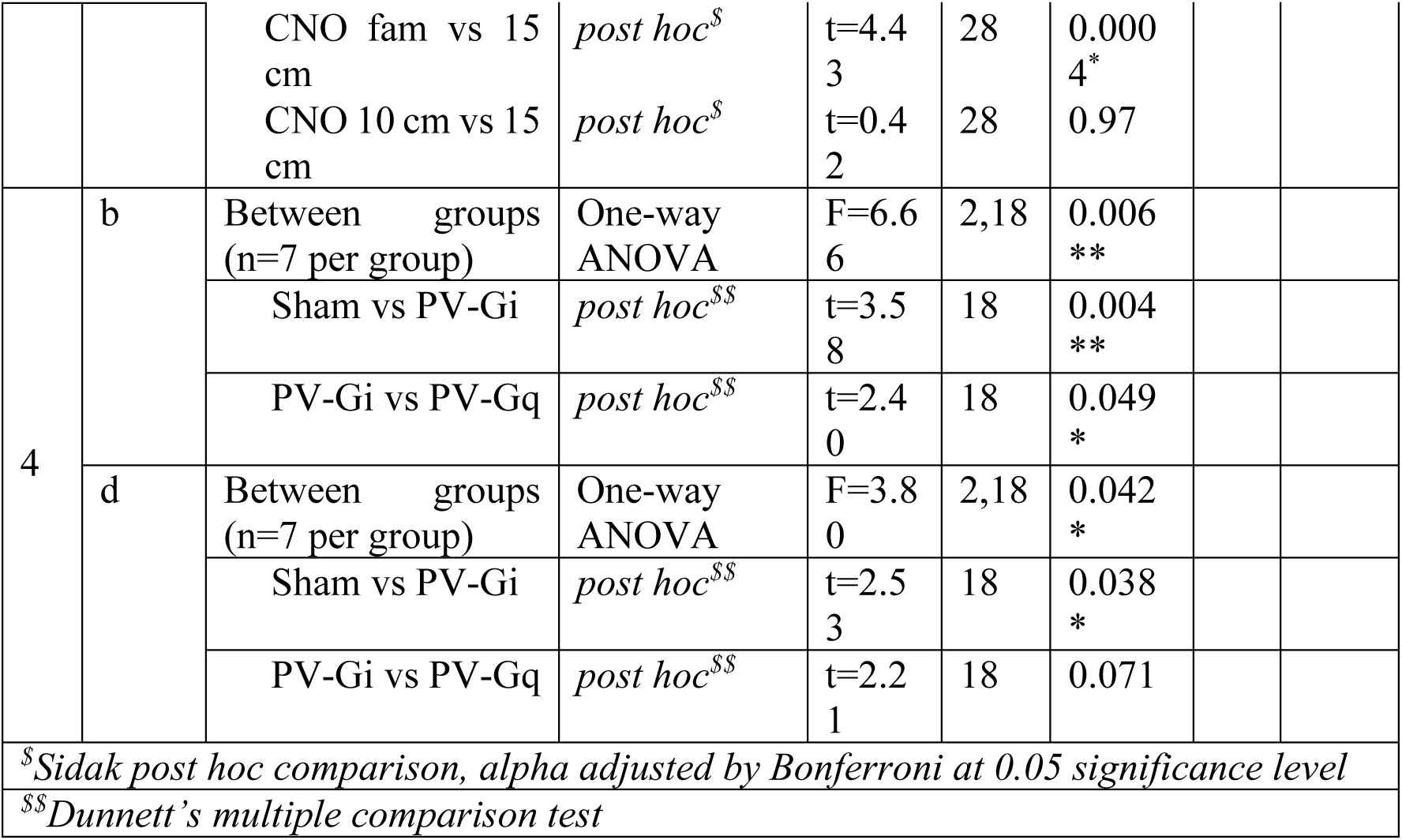
Statistical data corresponding to Fig. 1-2 and Fig. 4, including the statistical test, sample size (n), value of the statistic, degrees of freedom (df), associated *p* value, effect size (*η*^2^_*p*_) and power effect (1-β).

**Table S2.**
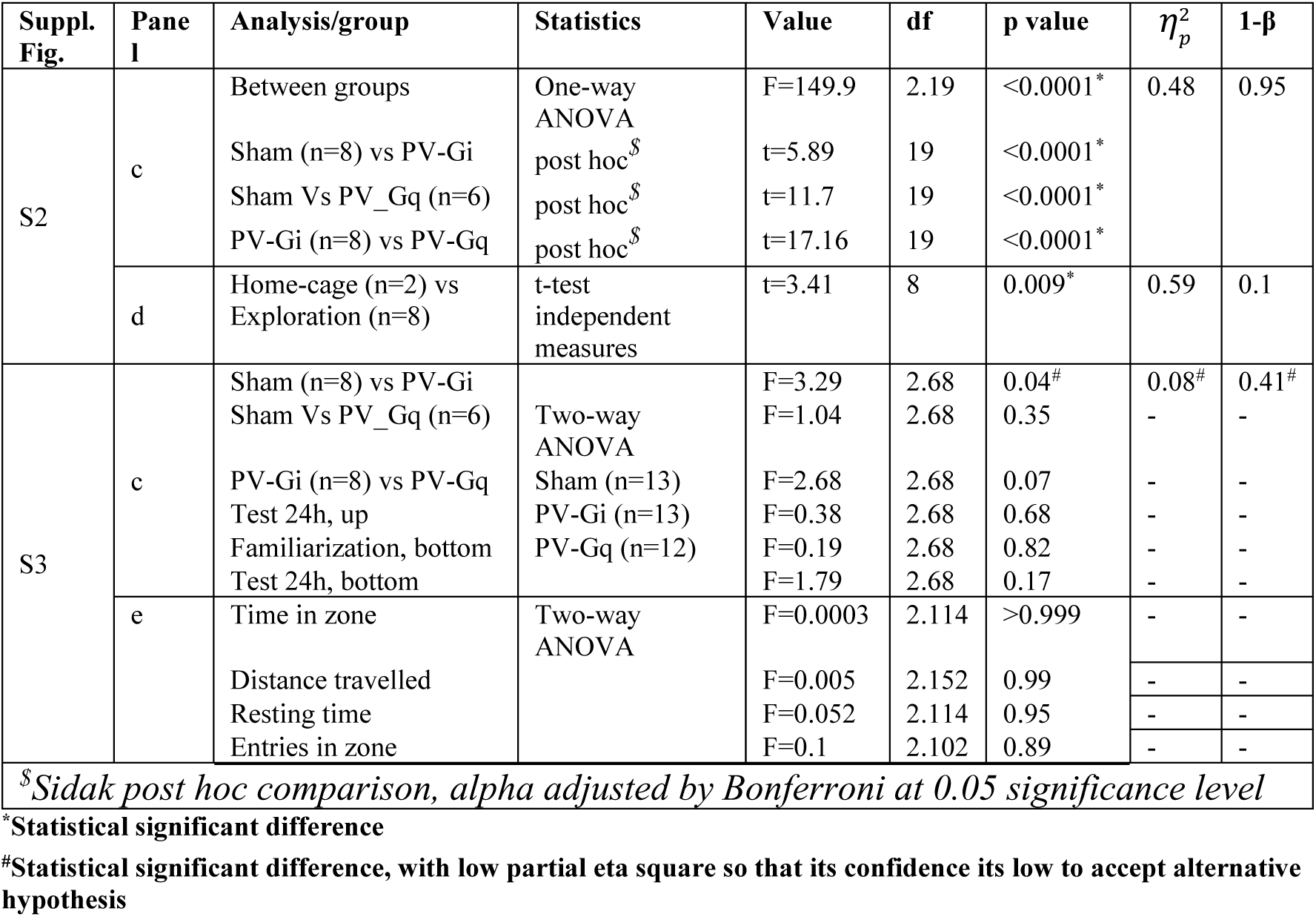
Statistical data corresponding to Supp. Fig. S2-S3, including the statistical test, sample size (n), value of the statistic, degrees of freedom (df), associated *p* value, effect size (*η*^2^_*p*_) and power effect (1-β).

